# Treatment of Huntington’s disease with a pan-HTT-targeting CRISPR nuclease

**DOI:** 10.64898/2025.12.20.695720

**Authors:** Katherine Tan, Daniela Del Bosque Siller, Alisha Y. Xiong, Anna X.A. Wang, Tristan X. McCallister, Sreya Mummadi, Luke A. St John, Tae Kyu Lee, Annika Carrillo, Daniel G. Renshaw, Richard H. Zhou, Colin K.W. Lim, Jack He, Christopher J. Fields, Michael R. Hayden, Thomas Gaj

**Affiliations:** Department of Bioengineering, Grainger College of Engineering, University of Illinois Urbana-Champaign, Urbana, IL 61801, USA; School of Molecular and Cellular Biology, University of Illinois Urbana-Champaign, Urbana, IL 61801, USA; High-Performance Biological Computing, Roy J. Carver Biotechnology Center, University of Illinois Urbana-Champaign, Urbana, IL 61801, USA; Centre for Molecular Medicine and Therapeutics, University of British Columbia, Vancouver, BC V5Z 4H4, CA; BC Children’s Hospital Research Institute, University of British Columbia, Vancouver, BC V5Z 4H4, CA; Department of Medical Genetics, University of British Columbia, Vancouver, BC V6T 1Z3, CA; Carl R. Woese Institute for Genomic Biology, University of Illinois Urbana-Champaign, Urbana, IL 61801, USA

## Abstract

Huntington’s disease (HD) is an inherited neurodegenerative disorder caused by an expansion of a CAG trinucleotide repeat in the huntingtin (HTT) gene, which leads to a mutant protein that destroys neurons in the brain. Despite intense effort, there remains no approved disease-modifying therapy for HD. Here we develop a pan-HTT-targeting CRISPR-Cas9 system that, when delivered to the striatum of R6/2 and YAC128 mice by AAV5, lowered mutant HTT mRNA and protein by 55-80% via its induction of frameshift-inducing indel mutations in HTT exon 1. Cas9 targeting improved motor coordination and locomotor activity, decreased anxiety-like deficits, reduced clasping and weight loss, limited striatal atrophy, and decreased the formation of intranuclear inclusions immunoreactive for the mutant HTT protein. In Hu21/21 mice, which carry the wild-type human HTT gene in lieu of the mouse ortholog, Cas9 lowered the HTT protein by 44% but induced no measurable behavioral deficits and had no adverse effect on neuronal viability, though its targeting was associated with neuroinflammation. Altogether, our results demonstrate the ability for a newly developed pan-HTT-targeting Cas9 system to affect HD-related phenotypes across models and provides insights into its tolerability.

## INTRODUCTION

Huntington’s disease (HD) is a dominantly inherited and fatal neurodegenerative disorder characterized by severe motor, cognitive and psychiatric symptoms^1^. Globally, HD affects approximately 5 individuals per 100,000 people but has the highest prevalence in North America, Europe and Australia^2,3^. In the United States, it’s estimated that more than 40,000 people are currently symptomatic for HD, with over 200,000 individuals believed to be at risk of developing the disorder. While medications can help to manage certain HD symptoms^4^, there remains no approved therapy that can alter the course of the disease, underscoring the need for new strategies to treat the disorder.

HD is caused by the expansion of a CAG trinucleotide repeat within exon 1 of the huntingtin (HTT) gene^5^. While unaffected individuals typically harbor between seven to 35 CAG repeats in the HTT gene, persons that are expected to develop HD within their lifetime carry 40 or more copies of it. This expansion leads to an elongated polyglutamine (polyQ) tract near the N-terminus of the HTT protein that increases its propensity to misfold and aggregate^6,7^, resulting in neurotoxicity^8^ and the gradual and progressive loss of neurons in the brain, with the striatum, a region involved in the control of movement and cognition, among the earliest affected^9^.

Though the precise pathological mechanism for how the mutant HTT (mHTT) protein causes HD remains uncertain, strategies capable of lowering it could offer a therapeutic benefit^10–12^. One modality with the ability to accomplish this is CRISPR-Cas9^13^, a versatile and efficient gene-editing technology^14,15^ with broad potential for gene therapy. CRISPR-Cas9 technology consists of two primary components: the Cas9 DNA endonuclease and a single guide RNA (sgRNA) that directs Cas9 to a genomic target via RNA-DNA base complementarity^13^. When bound to target DNA, Cas9 induces a double-strand break (DSB) that can stimulate non-homologous end-joining, an error-prone DNA repair pathway that rejoins the broken DNA ends but via a process that introduces insertion or deletion (indel) mutations at the cleavage site^16^. If introduced within a protein-coding region of a gene, a percentage of the indel mutations – specifically those whose length is not divisible by three – will alter the reading frame, an outcome that, in many cases, will create a downstream nonsense mutation that will be expected to lead to nonsense-mediated mRNA decay or premature translation termination, which will reduce the production of the encoded full-length protein^17,18^ (**Fig. 1A**).

**Figure 1.**
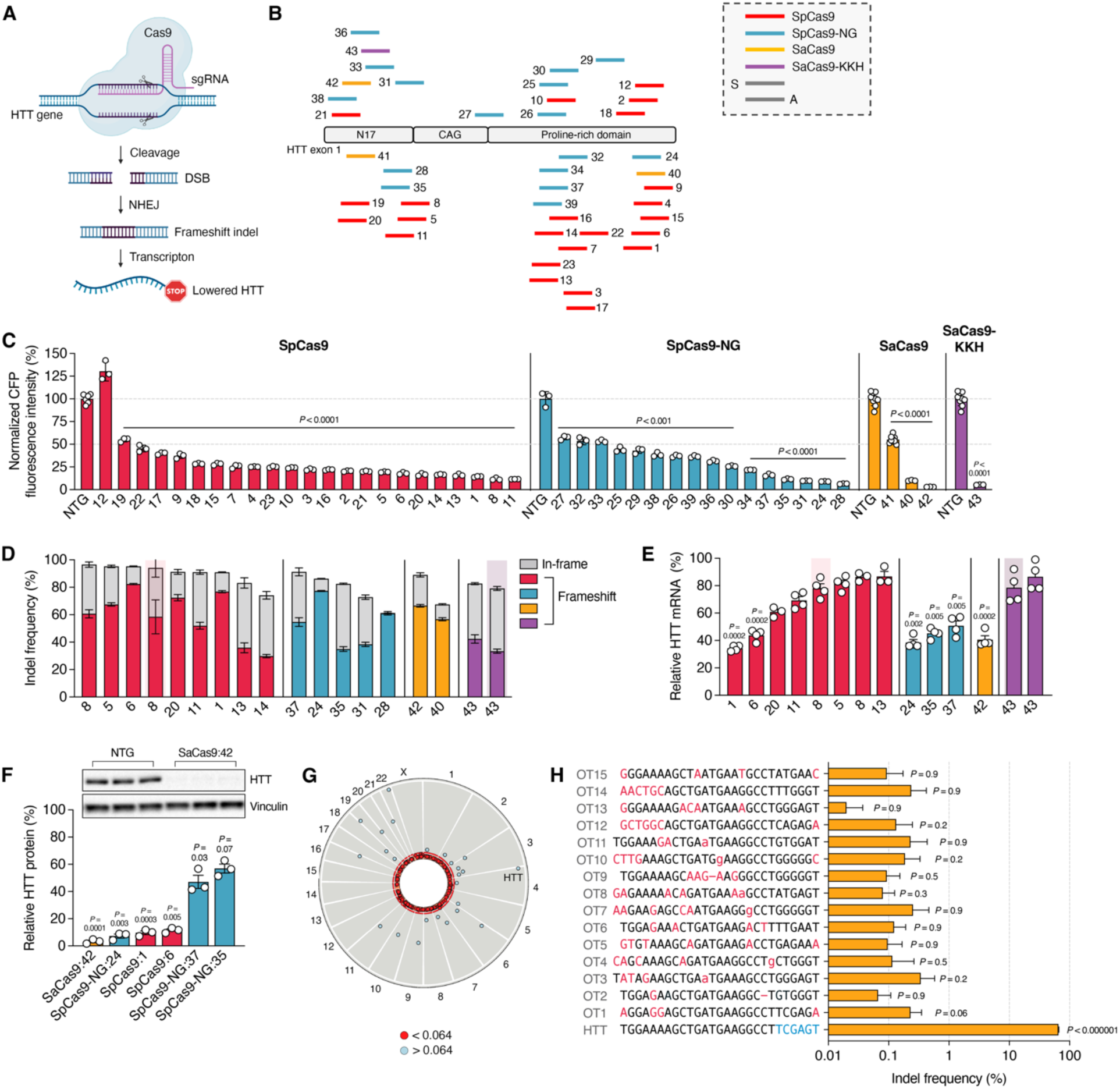
Cas9 targeting of HTT exon 1 reduced HTT mRNA and protein. (**A**) Cartoon of the mechanism for lowering HTT by CRISPR-Cas9. (**B**) Schematic of HTT exon 1 and the sgRNA binding sites for each Cas9 variant. S and A indicates whether sgRNA target sites are on the sense or antisense strands, respectively. (**C**) Cyan fluorescent protein (CFP) mean fluorescence intensity in HEK293T cells transfected with the HTT exon 1 reporter, the transactivator, and each Cas9:sgRNA variant. CFP for each Cas9:sgRNA variant was normalized to cells transfected with the same Cas9 variant with a non-targeting (NTG) sgRNA (n ≥ 3). (**D**) Indel frequencies in the HTT gene and (**E**) relative HTT mRNA in HEK293T cells transfected with each Cas9:sgRNA variant (n ≥ 3). (**D**) Red and purple shading denote the high-fidelity variants eSpCas9:8 and eSaCas9-KKH:43, respectively. (**E**) Data for each Cas9:sgRNA variant was normalized to cells transfected with the same Cas9 variant with a NTG sgRNA. (**F**) (Top) Western blot of the HTT protein and vinculin in HEK293T cells transfected with SaCas9:42 or SaCas9:NTG. (Bottom) Quantification of the HTT protein from HEK293T cells transfected with the indicated Cas9:sgRNA variants. Data for each Cas9:sgRNA variant was normalized to cells transfected with the same Cas9 variant with an NTG sgRNA (n = 3). (**G**) Modified Circos plot with the SaCas9:42 cleavage sites identified by Digenome-seq. The position of each circle relative to the inner ring indicates the normalized counts, with circles closer to the red inner ring signifying lower normalized counts. Normalized counts of >0.1 and <0.1 are blue and red circles, respectively. (**H**) Indel frequencies at the 15 highest-scoring sites for SaCas9:42 in HEK293T cells transfected with it (n = 3). All values indicate means and error bars indicate SD. Each Cas9:sgRNA variant was compared to their respective NTG using a one-tailed unpaired t-test, with exact *P* values shown. All data points are biologically independent samples.

We previously demonstrated the applicability of this approach for HD by showing that Cas9 could be programmed to target the HTT gene and facilitate the introduction of frameshift-inducing indels that lowered the HTT protein^19^. This was achieved via a non-allele-specific targeting strategy that was designed to lower both the mutant and wild-type HTT proteins but be applicable to a wider population than allele-specific approaches that rely on the presence of mHTT-associated single nucleotide polymorphisms^20–28^. However, though this Cas9-based system improved certain deficits in an HD rodent model, it was limited by a relatively low DNA editing efficiency^19^. Further, its tolerability and ability to improve HD-related deficits in an animal model that harbors the full-length (FL) HTT gene and more accurately recreates disease progression was undetermined. Given the potential benefits of a technology capable of permanently lowering the mHTT protein, we sought to optimize the capabilities of the Cas9-based system underlying this approach and to determine its ability to affect HD-related phenotypes across models, including one that could be used to analyze the tolerability of this pan-HTT-targeting strategy.

Here, we describe a new, highly efficient, pan-HTT-targeting CRISPR-Cas9 system for HD. By screening over 100 Cas9 variants programmed to target HTT exon 1, we identify a nuclease with the ability to decrease mHTT mRNA and protein by ∼55-80% when intrastriatally delivered to R6/2 and YAC128 mice. This Cas9 variant improved motor coordination and locomotor activity, decreased anxiety-like deficits, reduced clasping and weight loss, limited the volumetric loss of the striatum and decreased mHTT aggregation. To gain insight into the tolerability of this Cas9-based pan-HTT-targeting system, we intrastriatally delivered it to Hu21/21 mice, which carry two copies of the human wild-type HTT gene in lieu of the mouse ortholog. Cas9 targeting induced no apparent behavioral deficits, had no adverse effects on neuronal viability or neurite morphology and did not measurably affect striatal volume, though it increased neuroinflammation.

In summary, we developed a new pan-HTT-targeting CRISPR-Cas9 system capable of efficiently lowering the mHTT protein and demonstrate its ability to affect HD-related phenotypes across models. Our study also provides insights into its tolerability. Altogether, our results demonstrate the potential for CRISPR technology to treat HD.

## RESULTS

### Programming CRISPR-Cas9 to target HTT exon 1

We sought to develop a new, highly efficient, pan-HTT-targeting CRISPR-Cas9 system for HD. To target the HTT gene, we utilized the Cas9 nucleases from: (i) *Staphylococcus aureus* (SaCas9)^29^ whose relatively compact size can enable its delivery by a single adeno-associated virus (AAV) vector, and (ii) *Streptococcus pyogenes* (SpCas9)^13^ which, while necessitating two AAV vectors for delivery, can also mediate *in vivo* DNA editing^30^. In addition to these two enzymes, we also utilized: (iii) SaCas9-KKH^31^ and (iv) SpCas9-NG^32^, two engineered variants with increased protospacer adjacent motif (PAM) flexibility, which we expected would increase the number of targetable sites in HTT. Though Cas9 nucleases with further increased PAM flexibility have been developed, SaCas9-KKH and SpCas9-NG were among the most versatile variants available at the time this study was initiated.

As the N-terminal domain of the mHTT protein plays a role in the pathogenesis of HD^33^, we targeted exon 1 of the HTT gene, both upstream and downstream of the CAG repeat (**Fig. 1B**), reasoning that disrupting it would not only prevent the expression of the FL mutant protein and its downstream toxic effects, but also decrease the expression of HTT1a, a transcript variant that encodes the especially pathogenic mHTT exon 1 protein and is produced by a CAG-dependent alternative splicing event^34,35^. Using the CRISPR Guide RNA Design Tool, we identified 43 sgRNAs to target HTT exon 1, including three for SaCas9, which represented all of its possible sgRNAs, one for SaCas9-KKH, 23 for SpCas9, and 16 for SpCas9-NG. For SaCas9-KKH, SpCas9 and SpCas9-NG, sgRNAs were selected on the basis of minimum predicted specificity and efficiency scores (**Materials and Methods**).

To facilitate the identification of the most active Cas9:sgRNA variants, we utilized a previously implemented reporter that expresses exon 1 of the human HTT gene with 94 copies of the CAG repeat fused to cyan fluorescent protein (CFP) under the control of a tetracycline-responsive promoter element^36^ (**Fig. 1C**). Because this reporter links mHTT exon 1 expression to CFP, it can be used to measure Cas9-mediated disruption of the HTT gene via CFP fluorescence^19^.

To carry out our screen, we co-transfected human embryonic kidney (HEK) 293T cells with the HTT exon 1 reporter and a plasmid encoding the tetracycline-controlled transactivator to induce its expression, as well as an expression vector encoding one of the 43 Cas9:sgRNA variants. Flow cytometry was then used to measure CFP fluorescence at 72 hr post-transfection. In total, 37 of the 43 Cas9:sgRNA pairs reduced CFP fluorescence by at least 50% compared to their respective non-targeting variant (P < 0.01 for all; **Fig. 1C**), with 16 found to decrease it by at least 80% (P < 0.0001 for all 16; **Fig. 1C**), a number that we used as the cut-off for advancement to the second step of our screen. The most efficient variant, SaCas9:42, whose cleavage site is 23 base-pairs (bps) upstream from the start of the CAG repeat, decreased CFP fluorescence by ∼97% (P < 0.0001; **Fig. 1C**), a ∼50% improvement compared to the SaCas9 variant identified in our first study^19^, designated here as SaCas9:41.

Following the identification of the most efficient sgRNAs, we next determined their compatibility with high-fidelity (HF) versions of their respective Cas9 protein. These HF variants, which have a reduced propensity to cleave non-target DNA, included eSaCas9^37^ and SaCas9-HF^38^, eSpCas9(1.1)^37^, SpCas9-HF1^39^, SpCas9-HF2^39^, and SpCas9-HF4^39^. In the case of the PAM-flexible enzymes SaCas9-KKH and SpCas9-NG, the mutations that make up eSaCas9 and SaCas9-HF (for SaCas9-KKH) and eSpCas9(1.1), SpCas9-HF1, SpCas9-HF2, and SpCas9-HF4^39^ (for SpCas9-NG) were grafted onto the parental enzymes to create six new hybrid nucleases.

Surprisingly, we found that the majority of the HF variants had diminished activity compared to their native form when evaluated using the HTT exon 1 reporter (**Fig. S1**); however, two of the 58 nucleases were able to reduce CFP fluorescence by at least 80% (P < 0.001 for both; **Fig. S1**). These nucleases, eSpCas9(1.1):8 and eSaCas9-KKH:43, were advanced for additional testing with the 16 other Cas9:sgRNA variants that reduced CFP fluorescence by >80%.

As the next step of the screen, we determined the ability of the 18 Cas9:sgRNA variants identified above to target the endogenous HTT gene in HEK293T cells. Following their co-transfection with a plasmid that expressed a puromycin N-acetyl-transferase (PAC) gene, we conducted a puromycin selection to enrich for transfected cells, a method that enables a higher resolution analysis of Cas9-mediated outcomes^40^, which we reasoned would enable a more effective comparison between these active variants. We then used next-generation sequencing (NGS) to measure from the enriched cells the percentage of HTT exon 1 reads with indel mutations at each sgRNA target site, which is expected to be indicative of Cas9-mediated targeting.

Of the 18 variants tested, 14 induced indel mutations with frequencies exceeding 75% at six-days post-transfection (**Fig. 1D**), including seven sgRNAs for SpCas9 (8, 5, 6, 20, 11, 1, and 13; most active to least active), one sgRNA for eSpCas9(1.1) (8), three sgRNAs for SpCas9-NG (37, 24, and 35; most active to least active), one sgRNA for SaCas9 (42), one sgRNA for SaCas9-KKH (43), and one sgRNA for eSaCas9-KKH (43). Though the percentage of frameshift mutations varied based on the Cas9:sgRNA pair, ∼75%, ∼85%, and ∼89% of the indels for SaCas9:42, SpCas9:1, and SpCas9-NG:24, respectively, were frameshift mutations (**Fig. 1D**).

Using qPCR, we next evaluated the effect that the 14 most efficient Cas9:sgRNA variants had on HTT mRNA, which we measured at six-days post-transfection using validated primers that bind to the 5’ untranslated region (UTR) of the HTT transcript^41^. In total, six of the 14 variants decreased HTT mRNA by at least 50% (P < 0.01 for all six; **Fig. 1E**), including SaCas9:42, SpCas9-NG:24, and SpCas9:1, which reduced it by ∼60%, ∼62%, and ∼65%, respectively.

To examine if the Cas9:sgRNA pairs that most efficiently decreased HTT mRNA (SpCas9: 1; SpCas9-NG: 24, 35, and 37; and SaCas9:42) also lowered HTT protein, we conducted western blot on lysates from puromycin-enriched cells at 12 days post-transfection. Using the anti-HTT clone 1HU-4C8, which binds to an epitope between residues 181 and 810 of the HTT protein, we found that four of the six nucleases (SaCas9:42, SpCas9-NG:24, SpCas9:1, and SpCas9:6) decreased the FL protein by at least 80% relative to their respective controls (P < 0.01 for all; **Fig. 1F** and **Fig. S2**), with SaCas9:42 and SpCas9-NG:24 each found to decrease it by ∼90%. As the decrease in HTT protein for these variants exceeded their measured indel frequency, we hypothesize that the six additional days of culture, which we used to enable the turnover of the HTT protein, may have enabled the accumulation of additional indel mutations.

We next analyzed the genome-wide cleavage specificities of SaCas9:42 and SpCas9-NG:24, the SaCas9- and SpCas9-based systems that most effectively targeted HTT exon 1 and lowered HTT, as determined by a composite Activity Score (**Table S1**). Cleavage specificity was determined using Digenome-seq^42^, a method that first involves digesting purified genomic DNA with a Cas9 ribonucleoprotein (RNP) and then using whole-genome sequencing (WGS) to detect the cleaved fragments, which we measured as counts. In total, we identified 106 cleavage sites for SpCas9-NG:24, each of which had a normalized score of at least 0.064, a number we calculated as the ratio of counts for a given off-target site versus those for HTT (**Table S2** and **Fig. S3**). By comparison, we identified just 24 cleavage sites with normalized scores above 0.064 for SaCas9:42 (**Fig. 1G** and **Table S2**), though SaCas9:42 was found to have a larger percentage of sites with normalized scores below 0.01, which, based on the cleavage scores observed with SaCas9 in the absence of an sgRNA (**Table S2)**, we considered as likely background. As a positive control for this analysis, we conducted Digenome-seq with SpCas9 and a previously characterized sgRNA that targets the HBB gene^42^, which revealed many of the expected cleavage sites, including OT3, OT1, HBB_75 and HBB_48 (**Table S2**), reinforcing the validity of our methods.

Given it possessed the more favorable genome-wide cleavage profile, we next determined if SaCas9:42 modified any of the off-target cleavage sites identified by Digenome-seq in human cells. Using NGS, we found that none of the 15 highest scoring cleavage sites had indel mutations at a frequency higher than 0.25% in HEK293T cells transfected with SaCas9:42 (P > 0.05 for all; **Fig. 1H**). This analysis, which was conducted with no puromycin selection, showed that SaCas9:42 introduced indels in the HTT gene with a frequency of ∼63% (**Fig. 1H**).

Thus, on the basis of its ability to efficiently introduce frameshift-inducing indel mutations in HTT exon 1, which lowered HTT mRNA and protein, its favorable genome-wide cleavage profile, and its compact size, which can enable its *in vivo* delivery from a single AAV vector, SaCas9:42 was selected for further evaluation.

### AAV5-mediated delivery of SaCas9 to R6/2 mice reduced HTT mRNA and mutant protein inclusions and provided broader functional benefits than AAV1

We next evaluated the ability of SaCas9:42 to target HTT *in vivo* and improve deficits in HD mouse models. We first determined this in R6/2 mice^43^, which carry exon 1 of the human HTT gene with ∼150 CAG repeats and a ∼1 kb fragment of the HTT 5’ UTR that drives expression of the pathogenic fragment. R6/2 mice display progressive neurological abnormalities, including loss of coordination, irregular movements and weight loss, and develop intranuclear inclusions in neurons that are immunoreactive for mHTT.

Given the evidence to suggest that lowering total HTT protein in the striatum of adult animals may be more tolerated^44–48^ than brain-wide approaches^46,49,50^, we chose to locally deliver SaCas9 to the striatum using AAV. For our initial study, we compared delivery by AAV1 and AAV5, two AAV serotypes that can transduce neurons in the brain^51,52^ but demonstrate varying degrees of spread and retrograde transport^53^.

R6/2 mice develop an aggressive, early-onset phenotype, with most symptoms emerging by six to eight weeks of age. Given the need for an early intervention in this model, we intrastriatally dosed four-week-old R6/2 mice with ∼1.2 x 10^11^ vector genomes (VGs) of AAV1 or AAV5 encoding SaCas9 with sgRNA:42 (AAV1/5-SaCas9:HTT) or an sgRNA targeting the mouse Rosa26 safe-harbor locus (AAV1/5-SaCas9:mRosa26; **Fig. 2A**), which served as the negative control. Because the presence of the mHTT protein in neurons has been shown to be a driver of disease in R6/2 mice^54^, we used a minimal version of the neuron-specific human synapsin-1 (hSyn) promoter^55^ to express SaCas9 (**Fig. 2A**). Additionally, to facilitate the isolation of transduced nuclei to analyze SaCas9-mediated outcomes, we co-injected a subset of animals with an equal dose of a second AAV1 or AAV5 vector carrying a hSyn-driven EGFP variant fused to the KASH (Klarsicht/ANC-1/Syne-1 homology) domain^56^. Since this EGFP variant localizes to the outer nuclear membrane of cells, it can enable the isolation of transduced nuclei by fluorescence-activated cell sorting (FACS)^57,58^ (**Fig. 2A**), as a high proportion of cells are expected to be transduced by both vectors.

**Figure 2.**
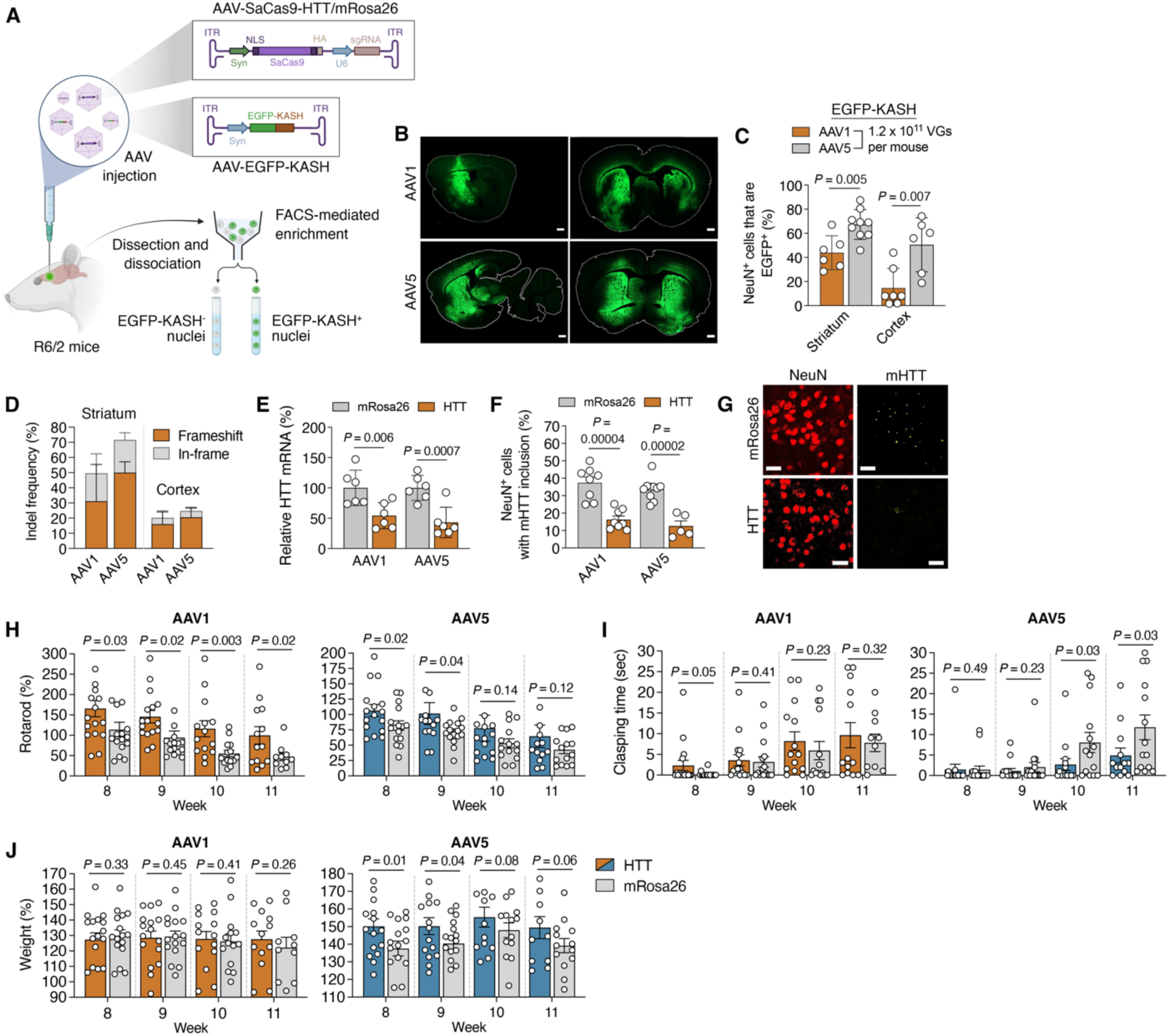
AAV5-mediated delivery of SaCas9 to R6/2 mice reduced mutant HTT mRNA and inclusions and provided broader functional benefits than AAV1 delivery. (**A**) Cartoon of the injection scheme in R6/2 mice and the plan to enrich EGFP-KASH^+^ cells by fluorescence-activated cell sorting (FACS). (**B**) Brain-wide EGFP-KASH fluorescence and (**C**) the percentage of NeuN^+^ cells that are positive for EGFP-KASH in the striatum and cortex (n ≥ 5) of R6/2 mice six-weeks after injection with 1.2 x 10^11^ vector genomes (VGs) each of AAV1/5-SaCas9:HTT/mRosa26 and AAV1/5-EGFP-KASH. >200 NeuN^+^ cells counted per region per mouse. Scale bars, 1 mm. (**D**) Indel frequencies (n = 4) and (**E**) relative HTT exon 1 mRNA (n = 6) in EGFP-KASH^+^ cells from R6/2 mice from **B**, **C**. (**E**) AAV1/5-SaCas9:HTT values were normalized to AAV1/5-SaCas9:mRosa26. (**F**) Percentage of NeuN^+^ cells with at least one EM48^+^ inclusion in the striatum of R6/2 mice from **B**, **C** (n ≥ 5) and (**G**) representative immunofluorescence from this analysis. >200 NeuN^+^ cells counted per mouse. Scale bars, 25 μm. (**D**, **E**, and **F**) Values indicate means and error bars indicate SD. (**H**) Rotarod, (**I**) clasping time and (**J**) weight of R6/2 mice injected with 1.2 x 10^11^ VGs of AAV1/5-SaCas9:HTT/mRosa26 (AAV1-HTT = 17; AAV1-mRosa26 = 16; AAV5-HTT = 16; AAV5-mRosa26 = 16). Weight and rotarod values for each mouse were normalized to their values at six weeks of age. (**H**, **I** and **J)** Values indicate means and error bars represent SEM. All comparisons conducted using a one-tailed unpaired t-test, with exact *P* values shown. All data points are biologically independent samples.

We first analyzed delivery to R6/2 mice at six weeks post-injection. Though EGFP-KASH was observed in the striatum and cortex of mice dosed with both AAV1 and AAV5 (**Fig. 2B**), an increased number of EGFP-KASH^+^ cells were measured in both areas for animals injected with AAV5. Specifically, ∼67% and ∼50% of the analyzed NeuN^+^ cells in the striatum and cortex, respectively, were positive for EGFP-KASH in mice dosed with AAV5 versus ∼43% and ∼14% for AAV1 (**Fig. 2C and Fig. S4**), though we observed greater variability in cortical areas with AAV5.

After verifying delivery, we next used FACS to isolate EGFP-KASH^+^ cells from the striatum and cortex of R6/2 mice. Using NGS to measure the frequency of indel mutations in HTT exon 1 from enriched cells, we found that mice dosed with AAV5- and AAV1-SaCas9:HTT had indels in ∼72% and ∼50% of their analyzed reads, respectively (**Fig. 2D**), with ∼70% and ∼63% of the indel products found to be frameshift mutations (**Fig. 2D**). For EGFP-KASH^+^ cells enriched from the cortex, indels were detected in ∼25% and ∼20% of the HTT exon 1 reads for AAV5- and AAV1-SaCas9:HTT, respectively, (**Fig. 2D**), with ∼83% and ∼79% of the products found to be frameshift variants (**Fig. 2D**).

Using qPCR, we next measured the effect that SaCas9 had on HTT exon 1 mRNA in R6/2 mice. For EGFP-KASH^+^ cells enriched from the striatum, we found that both AAV vectors decreased HTT mRNA by at least 45% (P < 0.01 for both; **Fig. 2E**), with animals dosed with AAV5-SaCas9:HTT found to have ∼63% less HTT mRNA. Though indels were measured in cells from the cortex, we were unable to isolate enough cells by FACS to reliably measure changes in HTT mRNA.

R6/2 mice develop inclusions that consist of the N-terminal domain of the mHTT protein. Because of the technical challenges associated with accurately quantifying the soluble and insoluble forms of this protein in striatal lysate from R6/2 mice^59^, we instead determined whether SaCas9 decreased inclusions immunoreactive for the mHTT protein, which can be readily detected as early as four weeks of age. Focusing on NeuN^+^ cells in the striatum and using an antibody (EM48) that preferentially recognizes the N-terminal domain of the mHTT protein^60^, we measured by immunofluorescence that R6/2 mice treated by AAV5-SaCas9:HTT and AAV1-SaCas9:HTT had a ∼63% and ∼57% decrease, respectively, in the number of NeuN^+^ cells with at least one EM48^+^ inclusion (P < 0.0001 for both; **Fig. 2F-2G**), indicating the ability for SaCas9 to lower the HTT exon 1 protein in R6/2 mice.

R6/2 mice show a progressive decline in motor function that coincides with weight loss and clasping behavior, a dystonic posturing of the limbs indicative of neurological dysfunction. To determine whether SaCas9 improved these deficits, we next measured rotarod performance, weight and clasping in R6/2 mice during the disease phase, which, for this study, was weeks 8, 9, 10, and 11.

Though mice injected with AAV1-SaCas9:HTT had improved rotarod function compared to their littermate controls (P < 0.05 for weeks 8, 9, 10, and 11; **Fig. 2H**), animals treated with this vector showed no improvements in clasping or weight during the disease phase (P > 0.05 for all measurements; **Fig. 2H-J**). R6/2 mice dosed with AAV5-SaCas9:HTT, however, showed a 23-77% improvement in latency to fall on the rotarod (P < 0.05 for weeks 8 and 9; **Fig. 2H**), a nearly three-fold reduction in clasping time (P < 0.05 for weeks 10 and 11; **Fig. 2I**) and a 5-10% increase in weight (P < 0.05 for weeks 8 and 9; **Fig. 2J**) compared to their littermate controls, indicating that AAV5-mediated delivery led to greater functional improvements than AAV1.

Thus, because of its more favorable distribution in the brain and the broader behavioral improvements enabled by it, AAV5 was chosen to deliver SaCas9 for our subsequent studies. Beyond the observed functional improvements, these results demonstrate that, when targeted to HTT exon 1 in R6/2 mice, SaCas9 can decrease HTT mRNA and reduce mHTT-immunopositive inclusions.

### SaCas9-mediated targeting of HTT exon 1 in YAC128 mice lowered mutant HTT mRNA and protein and improved functional deficits

We next determined the ability of SaCas9 to target HTT exon 1 and affect HD-like phenotypes in a slower progressing model of HD, namely YAC128 mice^61–63^, which encode the FL human mutant HTT protein with 128 CAG repeats. In addition to motor and cognitive abnormalities, YAC128 mice develop inclusions immunoreactive for the mHTT protein in neurons and exhibit age-dependent atrophy of the striatum^61^.

In contrast to our first study in R6/2 mice, which involved dosing animals at four weeks of age, we intrastriatally injected YAC128 mice with AAV5-SaCas9:HTT/mRosa26 at three months of age, a time point that was previously used to evaluate other mHTT-lowering approaches in this model^64,65^. In parallel, we co-injected a subset of animals with AAV5-EGFP-KASH to enable the isolation of transduced cells for a higher resolution of analysis of SaCas9-mediated outcomes.

Because astrocytes have been observed to contribute to disease progression in YAC128 mice^66^ and may play a role in the pathogenesis of HD^67^, we used a ubiquitous promoter to express SaCas9 (**Fig. 3A**), specifically a 582-bp version of the CAG promoter comprised of the early CMV enhancer element and the chicken beta-actin promoter^68^ that expresses in neurons^69^ and astrocytes^70^. Additionally, we injected mice with an increased dose of AAV5, specifically 6 x 10^11^ VGs instead of the 1.2 x 10^11^ VGs used in R6/2 mice, as 6 x 10^11^ VGs led to stronger and more widespread expression in the brain of YAC128 mice with the CAG promoter (**Fig. 3B**). In particular, for animals dosed with 6 x 10^11^ VGs, we observed that ∼72% and ∼50% of the analyzed NeuN^+^ cells in the striatum and cortex, respectively, were positive for EGFP-KASH (**Fig. 3C and Fig. S5**). Contrary to expectations, however, we observed limited EGFP-KASH expression in GFAP^+^ cells (**Fig. S6**).

**Figure 3.**
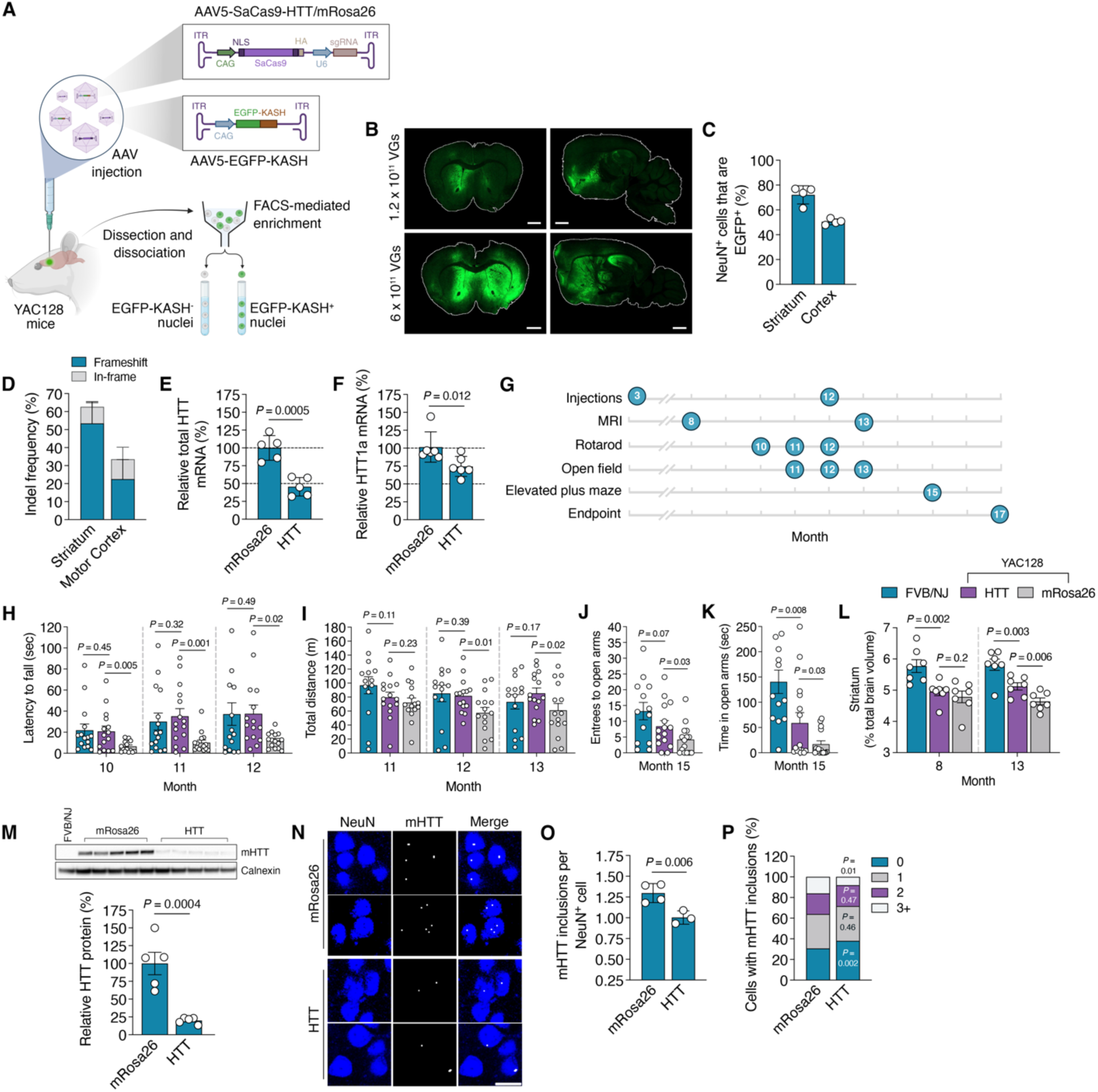
**SaCas9-mediated targeting of HTT exon 1 in YAC128 mice lowered mutant HTT mRNA and protein and improved functional deficits**. (**A**) Cartoon of the injection scheme in YAC128 mice and the plan to enrich EGFP-KASH^+^ cells by FACS. (**B**) Brain-wide EGFP-KASH fluorescence of YAC128 mice six-weeks after injection with 1.2 x 10^11^ or 6 x 10^11^ VGs of AAV5-EGFP-KASH. Scale bars, 1 mm. (**C**) The percentage of NeuN^+^ cells that are positive for EGFP-KASH in the striatum and cortex of YAC128 mice six-weeks after injection with 6 x 10^11^ VGs of AAV5-EGFP-KASH (n = 4). >270 NeuN^+^ cells counted per region per mouse. (**D**) Indel frequencies (n = 4), (**E**) total HTT mRNA (n = 5) and (**F**) HTT1a mRNA (n ≥ 5) in EGFP-KASH^+^ cells from the striatum of YAC128 mice two months after injection with 6 x 10^11^ VGs of AAV5-SaCas9:HTT/mRosa26 and 1.2 x 10^11^ VGs of AAV5-EGFP-KASH. (**E**, **F**) AAV5-SaCas9:HTT values were normalized to AAV5-SaCas9:mRosa26. (**D**, **E**, **F**) Values indicate means and error bars indicate SD. (**G**) Timeline for measurements. (**H**) Rotarod, (**I**) total distance traveled in an open field, (**J**) number of entrees to the open arms and (**K**) the time spent in the open arms for YAC128 mice injected with 6 x 10^11^ VGs of AAV5-SaCas9:HTT/mRosa26 at three months of age or uninjected FVB/NJ mice. (**H**) Rotarod values for each mouse were normalized to their values at three months of age (HTT = 15; mRosa26 = 15; FVB/NJ = 15 for **H** and **I**; FVB/NJ = 13 for **J** and **K**). (**H**, **I**, **J**, **K**) Values indicate means and error bars indicate SEM. (**L**) Volume of the striatum in 8- and 13-month-old YAC128 and FVB/NJ mice from **H**, **I**, **J**, **K** as a percentage of total brain volume (n = 7). (**M**) (top) Western blot of the full-length, soluble human mutant HTT protein and calnexin in striatal tissue from 17-month-old YAC128 and FVB/NJ mice from **H**, **I**, **J**, **K**. (Bottom) Quantification of western blot. Data for AAV5-SaCas9:HTT was normalized to AAV5-SaCas9:mRosa26 (n = 5). (**N**) Representative immunofluorescence staining for NeuN and mHTT via EM48 in the striatum of 17-month-old YAC128 mice from **H**, **I**, **J**, **K**. Scale bar, 15 μm. (**O**) Average number of EM48^+^ inclusions per NeuN^+^ cell from (**N**) and (**P**) the percentage of NeuN^+^ cells with 0, 1, 2, or >3 EM48^+^ inclusions (n ≥ 3). >156 cells counted per mouse. (**L**, **M**, **O**, **P**) Values indicate means and error bars indicate SD. All comparisons conducted using a one-tailed unpaired t-test, with exact *P* values shown. All data points are biologically independent samples.

Following delivery, we determined the ability of SaCas9 to target HTT. Using NGS to measure indel mutations in HTT exon 1 reads amplified from FACS-enriched EGFP-KASH^+^ cells, we found that YAC128 mice dosed with AAV5-SaCas9:HTT had indels in ∼63% and ∼36% of the reads from the striatum and cortex, respectively (**Fig. 3D**), with ∼85% of all products found to be frameshift variants (**Fig. 3D**). Consistent with the effect observed in R6/2 mice, EGFP-KASH^+^ cells from the striatum of mice dosed with AAV5-SaCas9:HTT had over two-fold less total HTT mRNA than the controls (P < 0.001; **Fig. 3E**), which we detected using primers that bind to the 5’ UTR of the HTT mRNA. Similar to our study in R6/2 mice, we were unable to isolate enough cells from the cortex by FACS to reliably measure HTT mRNA.

In addition to the FL HTT mRNA, YAC128 mice express HTT1a^34,71^, which encodes HTT exon 1, a potentially highly toxic form of the mutant protein that can be produced through a CAG-dependent alternative splicing event^34,35^. Using previously validated probes for HTT1a that bind upstream of the polyA2 site that contributes to its production^71^, we found that EGFP-KASH^+^ cells from the striatum of YAC128 mice injected with AAV5-SaCas9:HTT had ∼27% less HTT1a mRNA than the controls (P < 0.05; **Fig. 3F**), which we attribute to nonsense-mediated decay from Cas9 targeting.

YAC128 mice exhibit impaired motor coordination and reduced locomotor activity, as well as increased anxiety-like behavior, which can be observed by reduced exploration of open arms in an elevated plus maze. We next determined if SaCas9 improved these functional deficits, which we measured at different points from ten to 15 months of age (**Fig. 3G**), a period of time where these abnormalities are apparent.

Compared to the controls, YAC128 mice dosed with AAV5-SaCas9:HTT had both improved motor coordination (P < 0.01 for months 10 and 11; P < 0.05 for month 12; **Fig. 3H**) and locomotor activity (P < 0.05 for months 12 and 13; **Fig. 3I**), which was evidenced by up to a ∼175% improvement in latency to fall time on an accelerating rotarod and up to a ∼58% increase in total distance traveled in the open field, respectively. Notably, for both the rotarod and the open field, YAC128 mice treated with AAV5-SaCas9:HTT showed no difference compared to litter-matched wild-type FVB/NJ mice (P > 0.05 for all points for both assessments; **Fig. 3H and Fig. 3I**). 15-month-old YAC128 mice dosed with AAV5-SaCas9:HTT also showed nearly two-fold more entrees to the open arms of an elevated plus maze compared to controls (P < 0.05; **Fig. 3J**) and spent over twice the amount of time in the open arms (P < 0.05; **Fig. 3K**), showing no significant difference in the number of entrees to the open arms of the plus maze compared to wild-type FVB/NJ mice (P > 0.05; **Fig. 3J**), indicating that SaCas9 improved anxiety-like behavior. Though YAC128 mice exhibit increased weight gain compared to wild-type FVB/NJ mice, we measured no difference in weight between YAC128 mice injected with AAV5-SaCas9:HTT versus AAV5-SaCas9:mRosa26 (**Fig. S7**).

In addition to motor abnormalities and cognitive deficits, YAC128 mice show age-dependent atrophy of the striatum, which can be detected by magnetic resonance imaging (MRI)^72^. Though we observed no difference in striatal volume between treated and untreated eight-month-old YAC128 mice (P > 0.05; **Fig. 3L**), 13-month-old animals treated with AAV5-SaCas9:HTT were found to have ∼10% larger striatal volumes than those injected with AAV5-SaCas9:mRosa26 (P < 0.05; **Fig. 3L** and **Fig. S8**), though their volumes remained lower than wild-type FVB/NJ mice at this point (P < 0.05; **Fig. 3L**).

We next determined the effect that SaCas9 had on the mHTT protein, which we measured in 17-month-old animals. Based on a western blot using the anti-polyQ antibody IC2, which preferentially binds to the expanded polyQ repeat in HTT^73^, we measured that YAC128 mice injected with AAV5-SaCas9:HTT had ∼80% less FL soluble human mHTT in the striatum compared to the controls (P < 0.005; **Fig. 3M**). Using immunofluorescence, we also determined if SaCas9 affected mHTT aggregation, as YAC128 mice develop neuronal inclusions immunoreactive for mHTT. Within the striatum, 17-month-old animals dosed with AAV5-SaCas9:HTT were found to have ∼23% less EM48^+^ inclusions per NeuN^+^ cell (P < 0.01; **Fig. 3N-3O**). This included a ∼24% increase in NeuN^+^ cells with no detectable EM48^+^ inclusions and a ∼50% decrease in NeuN^+^ cells with three or more EM48^+^ inclusions (P < 0.05 for both; **Fig. 3P**), altogether indicating that SaCas9 decreased the formation of inclusions immunoreactive for the mHTT protein.

In sum, we find that SaCas9 can target HTT exon 1 in YAC128 mice, which we show decreased HTT mRNA and protein. SaCas9 also improved motor and cognitive deficits, limited the volumetric loss of the striatum, and lessened a neuropathological hallmark of HD in these mice.

### Tolerability of lowering wild-type HTT with Cas9 in Hu21/21 mice

Given the questions surrounding the tolerability of a pan-HTT-targeting strategy for HD, we next evaluated the effect(s) of using SaCas9 to lower the wild-type HTT protein in Hu21/21 mice^74^, a transgenic mouse model that carries two FL copies of the human HTT gene with 21 CAG repeats in lieu of the mouse wild-type HTT genes, which are knocked out. Hu21/21 mice carry an HTT exon 1 sequence that fully matches the human HTT reference sequence and have been used to analyze the tolerability of an AAV5-delivered pan-HTT-targeting microRNA that is the basis for AMT-130^75^.

As the mean age of onset for HD is typically between 35 to 44 years of age^76^, we chose to deliver SaCas9 to adult Hu21/21 mice – in this case six-month-old mice – which we intrastriatally dosed with 6 x 10^11^ VGs of AAV5-SaCas9:HTT/mRosa26, with the minimal version of the CAG promoter used to express SaCas9 (**Fig. 4A**). The ability of SaCas9 to target HTT was then evaluated by western blot and NGS, which revealed a ∼44% decrease in human HTT protein in striatal lysate from animals dosed with AAV5-SaCas9:HTT, which we probed using the anti-HTT clone 1HU-4C8 (P < 0.01; **Fig. 4B-4C**) and indels in ∼30% of the analyzed HTT exon 1 reads (**Fig. 4D**), respectively.

**Figure 4.**
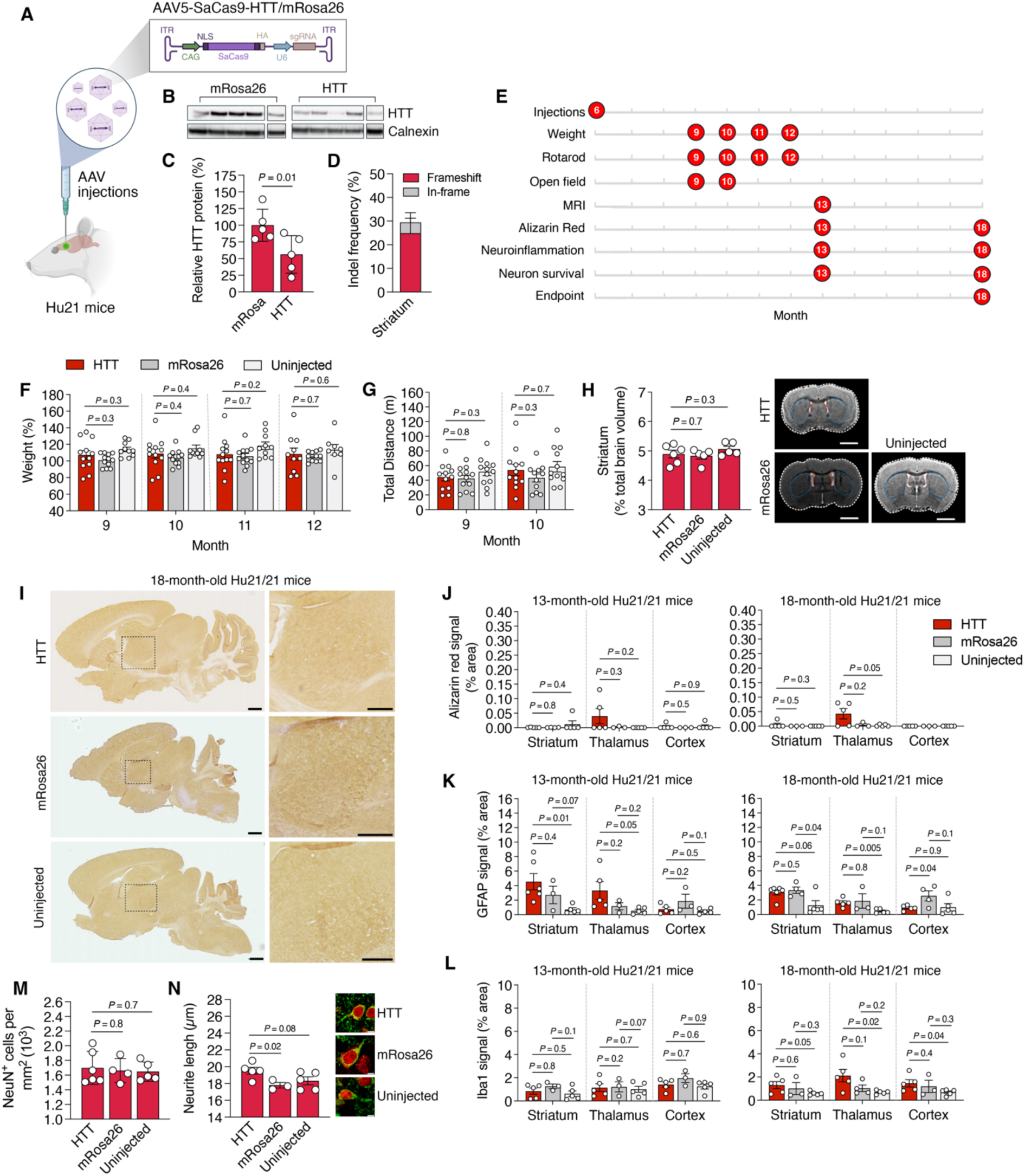
Tolerability analysis of AAV5-SaCas9:HTT in Hu21/21 mice. (**A**) Cartoon of the injection scheme in Hu21/21 mice. (**B**) Western blot of the full-length human HTT protein and calnexin in striatal tissue from Hu21/21 mice six months after injection with 6 x 10^11^ VGs of AAV5-SaCas9:HTT/mRosa26. (**C**) Quantification of western blot from **B**. Data for AAV5-SaCas9:HTT was normalized to AAV5-SaCas9:mRosa26 (n = 4). (**D**) Indel frequencies in the HTT gene amplified from 13-month-old Hu21/21 mice from **B** (n = 4). (**C**, **D**) Values indicate means and error bars indicate SD. (**E**) Timeline for measurements. (**F**) Weight and (**G**) total distance traveled in an open field for Hu21/21 mice injected with 6 x 10^11^ VGs of AAV5-SaCas9:HTT/mRosa26 at six months of age. Values from uninjected Hu21/21 littermates also shown. Weight values for each mouse were normalized to their values at six months of age (HTT = 12; mRosa26 = 12; uninjected = 10). (**F**, **G**). Values indicate means and error bars indicate SEM. (**H**) (Left) Volume of the striatum in 13-month-old Hu21/21 mice from **F, G** as a percentage of total brain volume. (Right) Representative images from the analysis (n ≥ 5). Blue and red lines denote the striatum and lateral ventricles, respectively. Scale bars, 2 mm. (**I**) Alizarin Red S staining of brain sections from 18-month-old Hu21/21 mice from **F, G**. Scale bars, 10 mm. Inset shows the thalamus for each section. Scale bars, 5 mm. ≥3 sections analyzed per animal. (**J, K, L**) Quantification of the area occupied by (**J**) Alizarin Red S, (**K**) GFAP, and (**L**) Iba1 in the striatum, thalamus and cortex of 13- and 18-month-old Hu21/21 mice from **F, G** (HTT = 5; mRosa26 = ≥3; uninjected = 5). (**M**) Quantification of the number of NeuN^+^ cells per mm^2^ from the striatum of 18-month-old Hu21/21 mice from **F, G** (HTT = 6; mRosa26 = 4; uninjected = 6). >250 NeuN^+^ cells counted per mouse. (**N**) (Left) Quantification of neurite length from Β-tubulin III^+^ cells in the striatum of 18-month-old Hu21/21 mice from **F, G**. (Right) Representative images. >24 Β-tubulin III^+^ cells traced per mouse. Scale bars, 5 μm. (HTT = 5; mRosa26 = 3; uninjected = 5). (**H**, **J**, **K**, **L**, **M**, **N**) Values indicate means and error bars indicate SEM. All comparisons conducted using a one-tailed unpaired t-test, with exact *P* values shown. All data points are biologically independent samples.

To determine whether lowering the HTT protein induced a functional deficit, we measured weight, locomotor activity and motor coordination in Hu21/21 mice from three to six months post-injection (**Fig. 4E**). At each point they were assessed, Hu21/21 mice injected with AAV5-SaCas9:HTT were found to have comparable weight (P > 0.05 for all; **Fig. 4F**) and locomotor activity (P > 0.05 for all; **Fig. 4G**) to the controls, which included Hu21/21 mice injected with AAV5-SaCas9:mRosa26 and an uninjected littermate cohort. In addition to demonstrating no appreciable difference in weight or locomotor activity, Hu21/21 mice dosed with AAV5-SaCas9:HTT also showed no difference in latency to fall times on an accelerating rotarod (P > 0.05 for all; **Fig. S9**), though it’s important to note that, potentially due to their increased weight, Hu21/21 mice display a decline in rotarod performance as early as four months of age^74^.

To determine whether lowering HTT induced structural deficits in the brain, we used MRI to measure striatal volume, observing that 13-month-old Hu21/21 mice dosed with AAV5-SaCas9:HTT had comparable volumes to both control groups (P > 0.05 for both; **Fig. 4H**), though mice dosed with AAV5-SaCas9:HTT and AAV5-SaCas9:mRosa26 had increased lateral ventricle volumes relative to the uninjected controls (P < 0.05 for both; **Fig. S10**), an effect that was previously observed in Hu21/21 mice following the high-dose injection of an AAV5 vector encoding a non-host transgene, specifically EGFP^75^.

The brain-wide elimination of mouse HTT by Cre-loxP recombination has been found to lead to calcium deposits in the thalamus^49,50^, which may be linked to the role the HTT protein plays in regulating and maintaining calcium homeostasis^77^. To determine if lowering HTT in Hu21/21 mice induced a similar abnormality, we analyzed the brains of 13- and 18-month-old mice with Alizarin Red S, a dye that binds to calcium ions and stains deposits orange-red.

While 13- and 18-month-old Hu21/21 mice dosed with AAV5-SaCas9:HTT had largely undetectable levels of Alizarin Red S in the striatum and cortex (P > 0.05 for all; **Fig. 4I-4J**), we observed increased signal in the thalamus, particularly in 18-month-old-animals, where, compared to the uninjected controls, Alizarin Red S levels were increased in total area by ∼16-fold (P = 0.054; **Fig. 4J**), though the area occupied by the signal remained low overall (AAV5-SaCas9:HTT: 0.043% ± 0.039%).

We next examined neuroinflammation. Consistent with a past study that observed astrogliosis in Hu21/21 mice following the high-dose delivery of a pan-HTT microRNA-encoding AAV5 vector^75^, we observed up to 6-fold more GFAP in the striatum and thalamus of 13-month-old mice injected with AAV5-SaCas9:HTT relative to the uninjected controls (P < 0.05 for striatum; P = 0.054 for the thalamus; **Fig. 4K** and **Fig. S11**). This increase was also evident in 18-month-old animals (P < 0.05 to uninjected controls; **Fig. 4K** and **Fig. S11**), though, at this point, GFAP was also elevated at similar levels in AAV5-SaCas9:mRosa26-injected mice, particularly within the striatum (P < 0.05 to uninjected control; **Fig. 4K** and **Fig. S11**).

Using Iba1, we also examined microgliosis. Though no increase was observed in 13-month-old mice dosed with AAV5-SaCas9:HTT, 18-month-old mice had significantly more Iba1 in the striatum, the thalamus and the cortex compared to the uninjected controls (P = 0.052 for striatum; P < 0.05 for thalamus and cortex; **Fig. 4L** and **Fig. S12**), though the percentage increased remained low overall (1-1.5%). No appreciable increase in Iba1 was observed for mice dosed with AAV5-SaCas9:mRosa26, suggesting microgliosis may be attributable to HTT targeting.

Last, we used immunofluorescence to analyze neuronal viability and morphology. Using the pan-neuronal marker NeuN and compared to the controls, we measured no difference in the number of NeuN^+^ cells per unit area in the striatum of 18-month-old Hu21/21 mice dosed AAV5-SaCas9:HTT (P > 0.05 for all; **Fig. 4M** and **Fig. S13**). Based on an immunofluorescent staining of the neurite marker β-tubulin III, we also measured no decrease in neurite length in 18-month-old mice injected with AAV5-SaCas9:HTT relative to the controls (**Fig. 4N**)

In sum, we find that lowering the HTT protein by nearly two-fold in Hu21/21 did not induce measurable deficits in weight, locomotor activity or motor coordination nor did it adversely affect striatal volume, neuronal viability or neurite morphology. However, increased neuroinflammation and a low percentage increase in calcium in the thalamus was observed in mice injected with AAV5-SaCas9:HTT.

## DISCUSSION

HD is a debilitating and invariably fatal neurodegenerative disorder with currently no approved disease-modifying therapies. We describe here a new, pan-HTT-targeting CRISPR-Cas9 system that can effectively lower HTT mRNA and protein by facilitating the introduction of frameshift-inducing indel mutations in the HTT gene and which has a favorable genome-wide cleavage profile. When intrastriatally delivered to R6/2 and YAC128 mice via AAV, this Cas9-based system decreased mutant HTT mRNA and protein by ∼55-80% and improved motor coordination, hindlimb clasping, and weight in R6/2 mice, and locomotor function, anxiety, and regional brain atrophy in YAC128 mice. While Cas9 was previously used to inactivate HTT through a similar non-allele-specific targeting strategy^19^, the Cas9 variant developed here is ∼50% more efficient than that variant.

The targeting strategy used here is non-allele-specific and thus expected to lower both the mutant and wild-type HTT proteins. Though the question of how safe this approach is remains debated, there is evidence to suggest that lowering total HTT in the striatum and cortical areas of adult animals^44–48^ may be more tolerated than strategies that aim to lower it throughout the whole brain^46,49,50^. To date, ASOs^65^, microRNAs^48,64,78,79^, small interfering RNAs^80–82^, and short hairpin RNAs^12^ have all been used to lower mHTT in a non-allele-specific manner, however, tominersen, a pan-HTT-targeting ASO developed by Ionis Pharmaceuticals and licensed to Roche^83^, failed to show a clinical benefit in a phase 3 trial and was determined to have an unfavorable risk/benefit profile^84^. Though it remains unknown whether this unfavorable profile was due to ASO-related side-effect(s) and/or from the nervous system-wide lowering of wild-type HTT, a post-hoc analysis revealed that a subset of patients, particularly younger individuals with less advanced disease, may have shown a benefit relative to older, more advanced patients. Given this observation, a new trial designed to evaluate tominersen in younger individuals at an earlier stage of disease with a lower dose of ASO and a less frequent dosing regimen has been initiated.

AMT-130, an AAV5-based gene therapy developed by uniQure that utilizes a pan-HTT-targeting microRNA^78,85–89^, is also under clinical evaluation. Unlike tominersen, which is administered intrathecally via a lumbar puncture and distributes throughout the nervous system, AMT-130 is surgically delivered to the caudate and putamen^88,90^, two major components of the striatum. Though the delivery of AMT-130 requires a more invasive and specialized procedure, it’s administered only once and directly to the striatum, a deep brain structure that can be challenging to access otherwise. In 2022, however, three of the 14 patients in the high-dose arm of the trial for AMT-130 experienced adverse reactions after its administration, which led to a halt of the study. Following a three-month pause, the trial for the high-dose group was resumed. More recently in September 2025, uniQure reported that a total of 12 trial precipitants that received a high-dose of AMT-130 had an apparent slowing in disease progression as determined by comparison to the Unified Huntington’s Disease Rating Scale and had less mean cerebrospinal fluid neurofilament light chain relative to baseline measurements, although this result was variable.

The HTT protein plays an important role in neuronal development^33^ and its knockout is embryonic lethal in mice^91,92^. Given the questions surrounding the tolerability of non-allele-specific approaches for HTT, we evaluated the effects of lowering the wild-type protein in adult Hu21/21 mice, which carry two copies of the FL human HTT gene with 21 CAG repeats in place of the mouse version of the gene. Our results show that lowering the HTT protein by ∼44% with Cas9 did not induce measurable deficits in weight, locomotor activity or motor coordination, nor did it adversely affect neuronal viability or neurite morphology. However, we did observe increased astrogliosis and microgliosis, as Hu21/21 mice dosed with AAV5-SaCas9:42 had up to a 4% increase in GFAP and a 1 to 1.5% increase in Iba1, with the striatum and thalamus generally found to the most affected. Increased calcium was also observed in the thalamus of Hu21/21 mice dosed with AAV5-SaCas9:42, though the area occupied by Alizarin Red S, the calcium dye used for this study, was low overall (0.043% ± 0.039%). It’s been shown that the brain-wide elimination of mouse HTT via Cre-loxP-mediated recombination can lead to the formation of severe calcium lesions in the thalamus^49,50^, which the original study investigators noted was reminiscent to those that develop in mouse models of Fahr disease and primary familial brain calcification^49^. It’s important to note, however, that the Alizarin Red S signal measured here was substantially lower than what was previously observed from the global elimination of mouse HTT^49,50^, though it’s possible that more extensive calcium lesions could have developed with additional time. It is also important to note that, though potentially serious, brain calcification can be a relatively common phenomenon. Up to 38% elderly persons have some brain calcification and its prevalence increases with age^93^, though its functional consequences in the context of HD will require further study.

The pan-HTT microRNA that is the basis for AMT-130 was also evaluated in Hu21/21 mice. While no clear anxiety-like abnormalities were observed in that study^75^, mice intrastriatally dosed with an AAV5 vector that carried the microRNA were found to exhibit an apparent dose-dependent decrease in latency to fall time on an accelerating rotarod and increased striatal atrophy^75^. Astrogliosis was also observed, but only in mice injected with the highest dose of the miRNA (1.3 x 10^11^ VGs per mouse) or with AAV5 vector carrying an EGFP transgene, which also induced microgliosis. While neuroinflammation was observed in our study, Hu21/21 mice dosed with AAV5-SaCas9:HTT exhibited no measurable deficits in weight, locomotor activity and rotarod function, and showed no evidence for striatal atrophy or neuronal loss. There are several important differences in the experimental design between the two studies that could help to explain these discrepancies. Beyond differences in dose and cargo, the study by Caron *et al*.^75^ delivered vector at an earlier time point (two months of age versus six months of age in our study) and conducted their analyses at different time points. Given the evidence that indicates that the developmental age at which HTT is lowered could influence the severity of the side effects from its depletion^46^, we hypothesize that the difference in the age of animals at dosing could be a contributing factor to the apparent discrepancies. Additionally, Caron *et al*.^75^ observed increased transduction in non-neuronal cells, which likely contributed to their apparent increased knockdown of HTT, which could also be a contributing factor. Unlike the study in Caron *et al*.^75^, we also note that our approach was not evaluated in a humanized HD model, such as Hu128/21^74^, which carry two full-length, human HTT transgenes and are heterozygous for an expanded CAG repeat. Evaluating the currently described Cas9-based strategy in such a model would provide insight into whether the net result of lowering both the mutant and wild-type HTT proteins is beneficial, though it’s important to note that the pan-HTT-targeting microRNA discussed above was also intrastriatally delivered to two-month-old Hu128/21 mice, where it was observed to improve psychiatric and cognitive phenotypes and prevent striatal atrophy, indicating an overall net benefit for pan-HTT-targeting.

It’s important to note that the Cas9-based strategy described here possesses several advantages compared to the antisense-based approaches previously used to lower HTT. First, because it edits DNA, Cas9 can permanently lower the mHTT protein and do so from only a single administration. This is in contrast to ASOs, which have a transient lifecycle and require re-dosing to sustain an effect. Second, cells that harbor Cas9-modified mHTT alleles with premature nonsense mutations are expected to produce no FL mHTT protein. This is in contrast to both ASOs and methods for RNA interference, which target HTT mRNA and thus may incompletely lower mHTT protein on a per cell basis. Thirdly, Cas9 is compatible with non-viral delivery methods^94^ and thus could enable hit-and-run editing if delivered as an mRNA or RNP. While AAV was used in this study and is a promising delivery vehicle, we nonetheless note that combining Cas9 with a non-viral delivery system could offer a means to improve tolerability. Additionally, it’s important to note that Cas9 has been used to modify the HTT gene in a number of ways^20,22,95–100^, including, intriguingly, by excising the CAG repeat^22,95–98^.

However, it’s worth noting that this approach could be confounded by competing DNA repair outcomes, as evidenced by a study in Hu97/18 mice, which found that only a relatively small percentage of the total edits for a dual sgRNA-based strategy to excise HTT exon 1 were the target deletion^101^, though this observation may be a model-dependent finding. Nonetheless, the expression of a second sgRNA poses a risk for increasing off-target effects. Beyond excising the CAG repeat, Cas9 has been used to inactivate^99^ and interfere^100^ with HTT expression in both allele-specific^20^ and non-allele-specific^99,100^ fashions. Compared to the Cas9-based approaches that aim to silence HTT with allele-specificity, we note that, since our approach does not rely on the presence of a mHTT-associated SNP to inactivate HTT, it’s expected to be applicable to the entire HD patient population.

Cas9, however, does have several weaknesses. First and foremost, it induces a DSB to initiate DNA editing. DSBs can activate DNA damage response pathways involving tumor suppressor proteins^102,103^ and carries a risk for inducing chromosomal translocations and rearrangements^104–106^. Cas9-induced DSBs can also capture AAV vectors, leading to their integration^107,108^. Though not analyzed here, these points will be important to assess in the future. Second, Cas9 relies on non-homologous end-joining to facilitate the introduction of the indel mutations that alter the HTT reading frame. Though our results show that ∼75-85% of the indels introduced by SaCas9:42 were frameshift variants, ∼15-25% of the indels were still in-frame and thus not expected to disrupt HTT expression. Third, Cas9, like other DNA editing technologies, can recognize off-target sites, which can lead to unintended mutations. Though Digenome-seq revealed a favorable specificity profile for SaCas9:42, with a subsequent targeted NGS analysis found to reveal none of the 15 highest scoring off-target sites identified by it were modified with frequencies greater than 0.25%, the continuous expression of Cas9 from an AAV vector can nonetheless lead to an accumulation of off-target effects over time. And fourth, Cas9 can be recognized as foreign by the immune system, which could elicit an immunogenic response that could trigger a number of different outcomes^109^, including potentially the loss of Cas9-modified cells. The implementation of strategies to help minimize the immunogenicity of both Cas9 and AAV will thus be important moving forward.

Over the past several years, many new CRISPR-based technologies have been developed to overcome of the limitations associated with traditional Cas9 nucleases^110^, with several of these technologies already used to target HTT. Most notably among these are base editors, which can create targeted point mutations without inducing a DSB. To date, base editors have been used to disrupt a key transcription factor binding site in the HTT promoter to affect the expression of the HTT gene^41^, induce exon skipping to prevent the proteolysis of the mHTT protein and inhibit the subsequent formation of mutant N-terminal fragments^111^, and convert the CAG repeat to a CAA repeat to prevent its further expansion^112,113^ as a means to potentially delay the onset of HD^114^. Another example of a recently emerged CRISPR technology that has been harnessed for HD is Cas13, an RNA-targeting modality that has now been used to lower HTT mRNA in both allele-specific^115,116^ and non-allele-specific fashions^69^. Though these technologies are all promising, our Cas9-based approach does offer certain advantages. First, compared to many base editing systems, the pan-HTT-targeting Cas9 variant developed here can fit within a single AAV vector, which is expected to make it more manufacturable and potentially more effectively delivered *in vivo*. Second, in contrast to base editors, whose deaminase domains can introduce non-specific modifications in RNA that could lead to mutations in proteins^117^, Cas9 has a limited risk for non-specifically cleaving RNA. And third, unlike RNA-targeting Cas13 systems, Cas9 does not require to be continuously expressed in cells to sustain an effect and does not collaterally cleave any nucleic acids after target cleavage^118^.

Finally, it’s important to note that a relatively high-dose of AAV5 vector was used in this study, particularly in YAC128 mice and Hu21/21 mice, which we used to ensure the efficient expression and distribution of Cas9. Given the concerns regarding potential toxicity of high-dose AAV gene therapy, future studies may be needed to identify an optimal compact promoter system capable of efficiently expressing Cas9 at lower doses in neuronal and non-neuronal cells.

In conclusion, we have developed a new, pan-HTT-targeting CRISPR-Cas9 system that can reduce HTT mRNA and protein and improve HD-related phenotypes across models. Our study also provides insights into the tolerability of this approach. These results altogether demonstrate the potential of CRISPR technology for HD.

## MATERIALS AND METHODS

### Plasmids

SaCas9, SaCas9-KKH and SaCas9-HF were expressed from pX601 (Addgene #61591), MSP1830 (Addgene #70708) and pCAG-CFP-SaCas9-HF (Addgene #134470), respectively. Prior to use, pX601 was digested with KpnI and NotI and ligated with the annealed oligonucleotides pX601-Stuffer-Fwd and pX601-Stuffer-Rev to remove the human U6 promoter and sgRNA.

eSaCas9 was created by Gibson Assembly from three PCR fragments amplified from pX601 using the primers: (i) eSaCas9-Fragment-1-Fwd and eSaCas9-Fragment-1-Rev, (ii) eSaCas9-Fragment-2-Fwd and eSaCas9-Fragment-2-Rev, and (iii) eSaCas9-Fragment-3-Fwd and eSaCas9-Fragment-3-Rev. The fragments were mixed at an equimolar ratio and inserted between the AgeI and EcoRI restriction sites of pX601 using the Gibson Assembly Master Mix (New England Biolabs; NEB) to generate pX601-eSaCas9.

To create the variants SaCas9-KKH-HF and eSaCas9-KKH, the point mutations that comprise SaCas9-HF and eSaCas9 were introduced into SaCas9-KKH-HF and eSaCas9-KKH by PCR using the primers: (i) SaCas9-KKH-HF-Fragment-1-Fwd and SaCas9-KKH-HF-Fragment-1-Rev, SaCas9-KKH-HF-Fragment-2-Fwd and SaCas9-KKH-HF-Fragment-2-Rev, SaCas9-KKH-HF-Fragment-3-Fwd and SaCas9-KKH-HF-Fragment-3-Rev, and SaCas9-KKH-HF-Fragment-4-Fwd and SaCas9-KKH-HF-Fragment-4-Rev for SaCas9-KKH-HF and (ii) eSaCas9-KKH-Fragment-1-Fwd and eSaCas9-KKH-Fragment-1-Rev, eSaCas9-KKH-Fragment-2-Fwd and eSaCas9-KKH-Fragment-2-Rev, eSaCas9-KKH-Fragment-3-Fwd and eSaCas9-KKH-Fragment-3-Rev for eSaCas9-KKH. MSP1830 was used as the template for the PCRs. The fragments for each variant were mixed at an equimolar ratio and then inserted into the AgeI and NotI restrictions sites of MSP1830 using the Gibson Assembly Master Mix.

SpCas9, SpCas9-NG and eSpCas9(1.1) were expressed from hCas9 (Addgene #41815), pX330-SpCas9-NG (Addgene #117919) and eSpCas9(1.1) (Addgene #71814), respectively. Prior to use, pX330-SpCas9-NG and eSpCas9(1.1) were digested with PciI and XbaI and ligated with the annealed oligonucleotides pX601-Stuffer-2-Fwd and pX601-Stuffer-2-Rev to remove the human U6 promoter and sgRNA.

SpCas9-HF1, SpCas9-HF2, and SpCas9-HF4 were expressed from VP12 (Addgene #72247), MSP2135 (Addgene #72248), and MSP2133 (Addgene #72249), respectively.

To create the variants eSpCas9-NG(1.1), SpCas9-NG-HF1, SpCas9-NG-HF2, and SpCas9-NG-HF4, the point mutations that comprise eSpCas9(1.1), SpCas9-HF1, SpCas9-HF2, and SpCas9-HF4 were introduced into SpCas9-NG by PCR using the primers: (i) eSpCas9-NG(1.1)-Fragment-1-Fwd and eSpCas9-NG(1.1)-Fragment-1-Rev, eSpCas9-NG(1.1)-Fragment-2-Fwd and eSpCas9-NG(1.1)-Fragment-2-Rev, eSpCas9-NG(1.1)-Fragment-3-Fwd and eSpCas9-NG(1.1)-Fragment-3-Rev, and eSpCas9-NG(1.1)-Fragment-4-Fwd and eSpCas9-NG(1.1)-Fragment-4-Rev for eSpCas9-NG(1.1), (ii) SpCas9-NG-HF1-Fragment-1-Fwd and SpCas9-NG-HF1-Fragment-1-Rev, SpCas9-NG-HF1-Fragment-2-Fwd and SpCas9-NG-HF1-Fragment-2-Rev, SpCas9-NG-HF1-Fragment-3-Fwd and SpCas9-NG-HF1-Fragment-3-Rev, SpCas9-NG-HF1-Fragment-4-Fwd and SpCas9-NG-HF1-Fragment-4-Rev and SpCas9-NG-HF1-Fragment-5-Fwd and SpCas9-NG-HF1-Fragment-5-Rev for SpCas9-NG-HF1, (iii) SpCas9-NG-HF2-Fragment-1-Fwd and SpCas9-NG-HF2-Fragment-1-Rev and SpCas9-NG-HF2-Fragment-2-Fwd and SpCas9-NG-HF2-Fragment-2-Rev for SpCas9-NG-HF2, and (iv) SpCas9-NG-HF4-Fragment-2-Fwd and SpCas9-NG-HF4-Fragment-2-Rev and SpCas9-NG-HF4-Fragment-2-Fwd and SpCas9-NG-HF4-Fragment-2-Rev for SpCas9-NG-HF4. pX330-SpCas9-NG was used as the template for all PCRs. The fragments encoding each variant were mixed at an equimolar ratio and then inserted into the AgeI and SacI restrictions sites of pX330-SpCas9-NG using the Gibson Assembly Master Mix. All primer sequences for constructing the Cas9 variants are provided in **Table S3**.

sgRNAs for HTT exon 1 were identified by the CRISPR Guide RNA Design Tool in Benchling. For SpCas9, all sgRNAs with specificity and efficiency scores greater than 30 were selected for testing. For SpCas9-NG and SaCas9-KKH, all sgRNAs with a specificity score greater than 30 and which were not cross-compatible with SpCas9 and SaCas9, respectively, were selected for testing.

sgRNAs for SaCas9, SaCas9-KKH, SaCas9-HF, and eSaCas9 were expressed from BPK2660 (Addgene #70709), while sgRNAs for SpCas9, SpCas9-NG, SpCas9-HF, SpCas9-HF1, SpCas9-HF2 and SpCas9-HF4 were expressed from pSP-sgRNA (Addgene #47108). Oligonucleotides encoding each sgRNA were incubated with T4 polynucleotide kinase (NEB) for 30 min at 37°C and then annealed at 95°C for 5 min. Oligonucleotides were then cooled to 4°C at a rate of -0.1°C per sec. The duplexed oligonucleotides were then ligated into the BsmBI and BbsI restriction sites of BPK2660 and pSP-sgRNA, respectively. All primer sequences for the sgRNAs are provided in **Table S4**.

The HTT exon 1 reporter and the tetracycline-controlled transactivator were expressed from pTreTight-HTT94Q-CFP (Addgene #23966) and tTA/TRE-mCherry, which was previously described^19^.

The plasmid pAAV-hSyn-SaCas9-sgRNA was previously described^19^. sgRNA:42 was amplified from BPK2660-sgRNA:42 using the primers sgRNA:42-Scaffold-Fwd and sgRNA:42-Scaffold-Rev. The ensuing amplicon was then inserted into the KpnI and NotI restriction sites of pAAV-hSyn-SaCas9-sgRNA to create pAAV-hSyn-SaCas9-sgRNA:42. sgRNA:mRosa26 was obtained as a gBlock (IDT) and amplified using the primers sgRNA:mRosa26-Scaffold-Fwd and sgRNA:mRosa26-Scaffold-Rev. The ensuing amplicon was then inserted into the KpnI and NotI restriction sites of pAAV-hSyn-SaCas9-sgRNA to create pAAV-hSyn-SaCas9-sgRNA:mRosa26.

To create the plasmids pAAV-CAG-SaCas9-sgRNA:42 and pAAV-CAG-SaCas9-sgRNA:mRosa26, the CAG promoter was first released from pAAV-CAG-C-Npu-SaPE2 by a restriction digestion using XbaI and AgeI and inserted into the same restriction sites of pAAV-hSyn-SaCas9-sgRNA:42 and pAAV-hSyn-SaCas9-sgRNA:mRosa26 to replace the hSyn promoter. All primer sequences for the pAAV plasmids are provided in **Table S3**.

Sanger sequencing (ACGT) was used to confirm the identity of each plasmid.

### Cell culture

HEK293T cells were cultured in Dulbecco’s modified Eagle’s medium (DMEM; Corning) supplemented with 10% (v/v) fetal bovine serum (FBS; Gibco) and 1% (v/v) antibiotic-antimycotic (Gibco) in a humidified 5% CO2 incubator at 37°C.

For experiments involving the HTT exon 1 reporter, HEK293T cells were seeded onto a 96-well plate at a density of 2 x 10^4^ cells per well and transfected the following day with 100 ng of pTreTight-Htt94Q-CFP, 100 ng of tTA/TRE-mCherry, 500 ng of Cas9 plasmid and 300 ng of sgRNA.

For experiments that targeted the human HTT gene, HEK293T cells were seeded onto a 24-well plate at a density of 2 x 10^5^ cells per well and transfected the following day with 600 ng of Cas9, 400 ng of sgRNA, and 100 ng of the PAC-encoding plasmid. Transfections were conducted with Lipofectamine 3000 according to manufacturer’s instructions.

### Flow cytometry

HEK293T cells were harvested 72 hours post-transfection, washed once with phosphate-buffered saline (PBS), and strained into single-cell suspensions using 5 mL round-bottom polystyrene test tubes with a 35 µm nylon mesh cell strainer snap cap (Falcon). CFP fluorescence was then measured using a BD LSR Fortessa Flow Cytometry Analyzer (Roy J. Carver Biotechnology Center Flow Cytometry Facility, University of Illinois Urbana-Champaign). A minimum of 50,000 events were recorded for each sample. Data was analyzed using the FACSDiva software (BD Biosciences).

### NGS

Genomic DNA was isolated from cells using the DNeasy Blood and Tissue Kit (Qiagen).

The sgRNA target sites in the human HTT gene were amplified using the KAPA2G Robust Hotstart PCR Kit (Roche) with 5% DMSO (v/v) and 400 nM each of one of three primer combinations: (i) HTT-NGS-Pair-1 for sgRNAs 42, 43, 20, 28, 8, 5, 35, 11, and 31; (ii) HTT-NGS-Pair-2 for sgRNAs 37, 13, and 14; and (iii) HTT-NGS-Pair-3 for sgRNAs 24, 6, 40, and 1. All primer sequences are provided in **Table S3**. PCR was conducted for 35 cycles using the following protocol: (i) initial denaturation at 95°C for 3 min; (ii) denaturation at 95°C for 20 s, annealing at 75°C, 63°C and 69°C for 15 s for primer pairs 1, 2 and 3, respectively, and extension at 72°C for 9 s, 11 s and 7 s for primer pairs 1, 2 and 3, respectively; (iii) final extension at 72°C for 30 s.

Barcoding for NGS was conducted by PCR using the KAPA2G Robust Hotstart PCR Kit with 3% DMSO (v/v) and 200 nM each of one of three different primer combinations for the sgRNA targets described above. PCR was conducted for 35 cycles using the following protocol: (i) initial denaturation at 95°C for 3 min; (ii) denaturation at 95°C for 20 s, annealing at 57.5°C for 15 s, and extension at 72°C for 11 s, 13 s and 9 s for primer pairs 1, 2 and 3, respectively; (iii) final extension at 72°C for 30 s.

Off-target sites for SaCas9:42 were amplified from HEK293T cells using the KAPA2G Robust Hotstart PCR Kit (Roche) as described by the manufacture. All primer sequences are provided in **Table S3**.

Following densitometry analysis by agarose gel electrophoresis, the PCR products were purified using a PureLink Quick Gel Extraction Kit (ThermoFisher Scientific) and then pooled and sequenced using a MiSeq (Illumina) from each end of the library using a Nano flow cell (Roy J. Carver Biotechnology Center, University of Illinois Urbana-Champaign). FASTQ files were created and demultiplexed using bcl2fastq v2.17.1.14 conversion software (Illumina).

CRISPResso was then used to quantify the frequency of indel mutations at each sgRNA site as described^119^.

### qPCR

RNA was isolated from cells using the PureLink RNA Mini Kit (ThermoFisher Scientific) and converted to cDNA using the iScript cDNA Synthesis Kit (BioRad) per the manufacturer’s instructions. qPCR was conducted on a 96-well plate with a QuantStudio 3 Real-Time PCR System (Thermo Fisher Scientific) using the iTaq Universal SYBR Green Supermix (BioRad). 50 ng of cDNA template was used in a reaction volume of 20 μL. qPCR measurements for each biological replicate were conducted at least in technical duplicates. HTT was detected using HTT-UTR-qPCR-Fwd and HTT-UTR-qPCR-Rev and compared to human GAPDH or mouse β-actin for each sample. The average fold-change was calculated using the 2ΔΔCT method.

Human HTT1a was measured using custom TaqMan probes (HTT-PolyA2-Fwd, HTT-PolyA2-Rev and HTT-PolyA2-Probe) with the TaqMan Fast Advanced Master Mix and was compared to mouse ATP5B (Mm01160389_g1; Thermo Fisher Scientific). Reaction volumes were 20 μL and probe concentrations were: 250 nM for HTT1a and 150nM for atp5b. TaqMan qPCRs were performed as recommended by the manufacturer’s instructions. Primer sequences are provided in **Table S3**.

### Western blot

Cells and tissue were lysed with radioimmunoprecipitation assay buffer (RIPA; 0.2% IGEPAL, 0.02% SDS, with Protease Inhibitor Cocktail [VWR] in PBS). Protein concentration was then determined using the DC Protein Assay Kit (BioRad) according to the manufacturer’s instructions.

Protein was electrophoresed by SDS-PAGE using 3 to 8% NuPAGE Tris-Acetate Mini Protein Gels (Thermo Fisher Scientific) in NuPAGE Tris-Acetate SDS Running Buffer (Thermo Fisher Scientific) with adapters for a Criterion Vertical Electrophoresis Cell (BioRad). Protein was electrophoretically transferred to a 0.45 μm polyvinylidene fluoride (PVDF) membrane using a Criterion Blotter (BioRad) in NuPAGE Transfer Buffer (Thermo Fisher Scientific) with NuPAGE Antioxidant (Thermo Fisher Scientific) and 20% methanol.

Following the transfer, membranes were blocked with 5% (v/v) Blotting-Grade Blocker (BioRad) in Tris-Buffered Saline (TBS; 10 mM Tris-HCl and 150 mM NaCl, pH 7.5) with 0.05% Tween 20 (TBS-T) for 1 hr. Membranes were then incubated with primary antibody in blocking solution overnight at 4°C. The following day, membranes were washed three times with TBS-T for 10 min and then incubated with secondary antibody in blocking solution for 1 hr at room temperature.

Membranes were then washed three times with TBS-T for 10 min and incubated with Super Signal West Dura Extended Duration Substrate (ThermoFisher Scientific). Chemiluminescence was measured using the ChemiDoc XRS+ System (Bio-Rad) and band intensities were quantified using Image Lab Software (Bio-Rad). Values were normalized to the reference protein for each sample.

The following primary antibodies were used: mouse anti-HTT clone 1HU-4C8 (1:1500 for HEK293T cells; 1:1000 for Hu21/21 mice; Millipore Sigma, MAB2166), mouse anti-polyQ clone 1C2 (1:4250; Millipore Sigma, MAB1574), rabbit anti-calnexin (1:5500; Millipore Sigma, C4731), and mouse anti-vinculin (1:1000; Proteintech, 66305-1-lg).

The following secondary antibodies were used: goat anti-rabbit horseradish peroxidase conjugate (1:4000; ThermoFisher Scientific, 31460) and goat anti-mouse horseradish peroxidase conjugate (1:2000; ThermoFisher Scientific, 31430).

### Digenome-seq

Genomic DNA was purified from HEK293T using the DNeasy Blood and Tissue Kit (Qiagen) and subsequently treated with RNase A (Thermo Fisher Scientific) for 10 min at 25°C. RNP complexes were then formed by incubating 100 nM of recombinant SaCas9 (IDT), SpCas9-NG (GenScript) and SpCas9 (IDT) proteins with 300 nM of sgRNA (IDT) for 10 min at 37°C. RNPs were then incubated with 10 μg of genomic DNA in the presence of NEBuffer 3.1 (100 mM NaCl, 50 mM Tris-HCl, 10 mM MgCl2, 100 μg/mL BSA, pH 7.9) for 8 hr at 37°C. Reaction mixtures were then treated with RNase A for 10 min at 25°C. RNP-digested genomic DNA was then purified using DNeasy Blood and Tissue Kit (Qiagen).

Shotgun genomic libraries was constructed by the Roy J. Carver Biotechnology Center (University of Illinois Urbana-Champaign). 200 ng of DNA per sample were sonicated using a Covaris ME220 (Covaris, MA) to an average fragment size of 400 bp. Libraries were then assembled with the Hyper Library Preparation Kit (Roche) using full-length barcoded adaptors without PCR amplification. Individually barcoded libraries were quantified by Qubit (Thermo Fisher Scientific) and run on a Fragment Analyzer (Agilent) to confirm the absence of adaptor dimers and to verify the expected size range of DNA. Libraries were then pooled in equimolar concentrations and quantified by qPCR on a CFX Connect Real-Time System (BioRad).

The barcoded libraries were loaded onto one S4 lane of a NovaSeq 6000 for cluster formation and sequencing (Illumina). Libraries were sequenced from both ends of the fragments, generating 150-bp reads from each end. FASTQ files were generated and demultiplexed using the bcl2fastq v2.20 Conversion Software (Illumina). Sequencing was performed by the Roy J. Carver Biotechnology Center (University of Illinois Urbana-Champaign).

The resulting reads were downloaded and analyzed on the IGB Biocluster using a Nextflow-based workflow. Reads were aligned to the human reference genome (GRCh37.p13) using bwa mem v0.7.17^120^, followed by coordinate-based sorting and indexing of aligned reads using SAMtools v1.12^121^. Alignments were then evaluated using Qualimap v2.2.1^122^ and Picard v2.10.1 (http://broadinstitute.github.io/picard/). Conversion of the aligned reads to a simple quantitative bigWig format^123^ was performed using deepTools v3.2.1^124^ with a standard genome browser. The resulting alignments were then processed using the standalone Digenome-seq tool^125^

### Injections

All animal procedures were approved by the Institutional Animal Care and Use Committee (IACUC) at the University of Illinois Urbana-Champaign and conducted in accordance with the National Institutes of Health (NIH) Guide for the Care and Use of Laboratory Animals. R6/2 mice (B6CBA-Tg(HDexon1)62Gpb/3J; Strain #006494) and YAC128 mice (FVB-Tg(YAC128)53Hay/J; Strain #004938) were obtained from the Jackson Laboratory. Hu21/21 mice were provided by the laboratory of M.R.H. All mice were housed in humidity and temperature-controlled rooms with a 12:12 hr light:dark cycle.

Mice were bred using a triad breeding scheme consisting of two wild-type females (B6CBAF1/J, Strain #100011 for R6/2 mice; FVB/NJ, Strain #001800 for YAC128 and Hu21/21 mice) and one transgenic male. Genotype was determined by PCR using genomic DNA extracted from an ear clip using the DNeasy Blood and Tissue Kit (Qiagen).

R6/2, YAC128 and Hu21/21 mice were injected using a drill and microinjection robot (NeuroStar) at stereotaxic coordinates anterior-posterior (AP) = 0.5 mm; medial-lateral (ML) = ±1.65 mm; and dorsal-ventral (DV) = -2.5 to -3.5 mm. 2 × 10^10^ VGs was delivered per site in 2.4 μL of saline for a total of 6 × 10^10^ VGs per hemisphere for R6/2 mice. 1 × 10^11^ VGs was delivered per site in 2.4 μL of saline for a total of 3 × 10^11^ VGs per hemisphere for YAC128 and Hu21/21 mice. AAV1 and AAV5 vectors were manufactured by the former University of Pennsylvania Vector Core.

All experimental groups were sex and litter balanced.

### Immunofluorescence

Mice were anesthetized using 3% isoflurane delivered through vaporizer in a closed chamber and transcardially perfused using PBS. Brain tissue was harvested and fixed for 48 hr in 4% paraformaldehyde (PFA) in PBS at 4°C before an incubation with 30% sucrose for 72 hr at 4°C. Brain tissue was sliced into 40 μm sagittal and coronal sections using a CM3050 S cryostat (Leica) and preserved in cryoprotectant (25% ethylene glycol, 25% glycerol, 50% 0.1M PBS pH 7.2) at -20°C until use.

For immunostaining, sections were washed three times with PBS and incubated with blocking solution (PBS with 10% [v/v] donkey serum [Abcam] and 1% Triton X-100) for 2 hr at room temperature. Primary antibodies were diluted in blocking solution and incubated with sections for 72 hr at 4°C. The sections were then washed three times with PBS and incubated secondary antibodies in blocking solution for 2 hr at room temperature.

The stained sections were then washed three times in PBS, mounted onto slides using VectaShield Hard Set Antifade Mounting Medium (Vector Laboratories) and visualized using a TCS SP8 confocal microscope (Leica; Beckman Institute Imaging Technology Microscopy Suite, University of Illinois Urbana-Champaign).

All image analyses were performed using Fiji. Neurite tracing and quantification were conducted using the Neuroanatomy plugin, with visible projections manually traced through a z-stack. Neurite lengths were measured for at least six neurons per tissue section and averaged per tissue section, with at least four sections analyzed per animal. The percentage area occupied by GFAP and Iba1 was measured using the threshold function in Fiji. The percentage of NeuN^+^ cells that were EGFP-KASH+, the number of NeuN^+^ cells per unit area, and the percentage of NeuN^+^ cells with mHTT inclusions was determined using the Cell Counter plugin in Fiji.

The following primary antibodies were used: mouse anti-HTT clone mEM48 (1:50; Millipore Sigma, MAB5374), rabbit anti-NeuN (1:500; Abcam, ab177487), chicken anti-NeuN (1:250; Millipore Sigma, ABN91), mouse anti-β-tubulin III (1:250; Millipore Sigma, T8578-200UL), chicken anti-GFP (1:500; Abcam, ab13970), rabbit anti-Iba1 (1:500; Wako Chemicals, 019-19741), and chicken anti-GFAP (1:1000; Abcam, ab4674).

The following secondary antibodies were used: donkey anti-chicken DyLight405 (1:150; Jackson ImmunoResearch, 705-165-155), donkey anti-rabbit DyLight405 (1:150, Abcam, ab175651), donkey anti-mouse Cy3 (1:150; Jackson ImmunoResearch, 715-165-150), donkey anti-chicken Cy3 (1:150; Jackson ImmunoResearch, 703-165-155), and donkey anti-rabbit Alexa647 (1:150; Jackson ImmunoResearch, 711-605-152).

### Neuronal nuclei isolation

Neuronal nuclei were isolated from dissected brain by FACS as described^58^. Briefly, following a transcardial perfusion with PBS, tissue from the cortex and striatum of mice were harvested and homogenized using a KIMBLE Dounce Tissue Grinder (Sigma-Aldrich) and incubated with 2 mL of Nuclei EZ Lysis Buffer (Sigma Aldrich) for 5 min. Homogenized tissue was then centrifuged at 500*g* for 10 min, washed once with Nuclei Suspension Buffer with 0.01% BSA, centrifuged again at 500*g* for 10 min and resuspended in 1 mL of Nuclei Suspension Buffer with 0.01% BSA. Suspensions were strained using a 5 mL round-bottom polystyrene tubes with a 35 µm nylon mesh cell strainer snap cap (Falcon) and then sorted using the Bigfoot Spectral Cell Sorter (Thermo Fisher Scientific; Roy J. Carver Biotechnology Center, University of Illinois Urbana-Champaign). At least 15,000 nuclei were collected per striatum and at least 3,000 nuclei were collected per cortex.

### Behavior

All measurements were conducted by a blinded investigator. Motor coordination was measured using a Rotamex-5 rotarod (Columbus Instruments) that accelerated from 4 to 40 rpm over 180 sec. The latency to fall was recorded for each mouse, with each session for each mouse comprised of five independent trials.

Clasping was measured by suspending mice by their tails for 30 sec. The amount of time that each mouse clasped their hindlimbs was recorded.

Locomotor function was measured with an open field. Mice were placed in a 17” x 17” square open arena (Panlab) for 10 min, with their total distance traveled tracked by the SMART Video Tracking System (Panlab).

Anxiety-like behavior was measured with an elevated plus maze. Mice were placed in the standard apparatus (Panlab) for 5 min, with their time spent in the closed versus open arms tracked by the SMART Video Tracking System (Panlab).

The weights of each mouse were recorded using an electronic scale.

### MRI

MRI scans were conducted using a 9.4T preclinical MRI system (BioSpec 94/30 USR, Bruker BioSpin MRI, Billerica) equipped with a gradient shim insert (B-GA12S HP) coil for mouse imaging. A four-channel mouse brain surface array coil was utilized to acquire high-resolution anatomical images of the brain. Animals were anesthetized with isoflurane (5% for induction, 1-2% for maintenance, mixed with 100% oxygen) delivered at 1 L per min. A three-dimensional T2-weighted fast spin echo sequence was used to acquire high resolution images of the mouse brain for volumetric characterization of the brain and its substructures.

Post-processing of MRI images was conducted using 3D Slicer v5.2.2 software (3D Slicer). Volumetric analysis of the brain and striatum were obtained using image registration and voxel-based morphometric techniques.

### Histochemistry

Following their removal from cryoprotectant, sections were washed three times with PBS for 5 min on a rocking shaker before and transferred onto microscope slides to air dry for at least 30 min at room temperature or until fully dry. Coverslips were then mounted using Permount Mounting Media (Electron Microscopy Sciences). Slides were then dipped into deionized water for 30 sec and incubated for 2 min in a 2% Alizarin Red (Sigma Aldrich) solution (pH 4.2) with 10% ammonium hydroxide. Slides were then rinsed gently with deionized water to remove excess solution and then dipped twenty times in 100% acetone. Slides were gently wiped and then dipped twenty times into a 1:1 mixture of acetone and xylene and then incubated with 100% xylene for 1 min and dried overnight at room temperature in a fume hood. Slides were stored at 4°C until use.

Slides were imaged using the NanoZoomer 2.0-HT Slide Scanner (Hamamatsu; Carl R. Woese Institute for Genomic Biology, University of Illinois Urbana-Champaign). Image analysis was performed using Fiji, with regions of interests first identified by the Allen Mouse Brain Atlas and subsequently segmented for a color threshold analysis in Fiji using the following default settings: saturation: 128-255, brightness: 192-255, and hue: 15-237 excluded (threshold settings were subject to adjustment based on background). From this, a mask was created for each segment whose percentage pixels was measured.

### Statistical analysis

Statistical analysis was performed using Prism 9.1 (GraphPad Software). For *in vitro* and *in vivo* studies, groups were compared using unpaired t-tests. For *in vitro* experiments, a minimum of three independent biological replicates were used. No statistical method was used to determine sample size for *in vitro* experiments. *For in vivo* experiments, the expected effect and error were informed from published literature, with the sample size determined by power calculations using α = 0.05 and β = 0.80. The sample size is the number of independent biological replicates, with all animals randomized into sex and litter-matching groups. For all qPCR measurements, if no amplification or minimal amplification of housekeeping genes was observed, the sample was removed from the analysis. All behavior assessments were conducted by a blinded investigator.

## DATA AVAILABILTIY

All experimental data are available from the corresponding author upon request.

## ACKNOWLEDGMENTS

We thank A. Hernandez for helpful discussion and assistance with library preparations for NGS and Anastasia Kuzmin for helping with animal breeding. This work was supported by a sponsored research agreement with Sarepta Therapeutics. Additional support for this work came from the NIH/NINDS (1U01NS122102-01A1 and 1R01NS123556-01A1 to T.G.), the NIH/NIGMS (5R01GM141296 to T.G.), the Muscular Dystrophy Association (MDA602798 to T.G.), the Simons Foundation (887187 to T.G.) and the Parkinson’s Foundation (PF-IMP-1950 to T.G.). A.Y.X was supported by a MCB Summer Undergraduate Research Fellowship. Cartoons were created with BioRender.

## AUTHOR CONTRIBUTIONS

T.G. conceived of the study. K.T. cloned the plasmids; K.T., D.D.B.S, A.Y.X., S.M. and L.S.J conducted cell culture; K.T. performed flow cytometry; K.T. and D.D.B.S. conducted NGS; K.T., A.Y.X. and C.K.W.L. conducted western blots; K.T., D.D.B.S, A.Y.X. and T.X.M. conducted qPCRs; KT., R.H.Z. and C.J.F. conducted Digenome-seq; K.T., D.D.B.S., A.Y.X., S.M., L.S.J., T.X.M. and C.K.W.L. conducted injections, K.T. and D.D.B.S. conducted FACS; D.D.B.S., A.Y.X, S.M., L.S.J, A.C., D.G.R., T.K.L and R.H.Z. conducted behavior; K.T., D.D.B.S., A.Y.X. and T.K.L conducted MRI. K.T., D.D.B.S., S.M., L.S.J., T.K.L., A.X.A.W., T.X.M. and J.H. conducted immunofluorescence; D.D.B.S and A.X.A.W. conducted histology; K.T., D.D.B.S., A.X.A.W., T.X.M., and J.H. conducted image analyses. M.R.H. provided mouse lines and critically evaluated and edited the manuscript; T.G, K.T, D.D.B.S., A.Y.X., T.X.M. and A.X.A.W. analyzed the data. T.G. wrote the manuscript with input from authors.

## DECLARATION OF INTERESTS

This work was supported by a sponsored research agreement with Sarepta Therapeutics. M.R.H. is the Chief Executive Officer of Prilenia Therapeutics and serves on the public boards of Ionis Pharmaceuticals, AbCellera, and 89bio.

